# The social, decoupled self: interpersonal synchronization of breathing alters intrapersonal cardiorespiratory coupling

**DOI:** 10.1101/2025.09.29.679159

**Authors:** Ivana Konvalinka, Natalie Sebanz, Guenther Knoblich

## Abstract

People synchronize their behavioural and physiological rhythms with each other during social interaction. While interpersonal synchronization has largely been associated with positive effects such as social bonding, some evidence suggests that it may also impair self-regulation and disrupt intrapersonal coordination. Because respiration and heart rhythms are weakly coupled within individuals, we investigated whether synchronizing breathing with another person alters intra-personal cardiorespiratory coupling. Across two experiments, participants synchronized their breathing reciprocally (bidirectional interaction), unidirectionally with a confederate, or with pre-recorded breathing signals, while respiration and electrocardiography (ECG) were continuously measured. Relative phase analyses revealed that bidirectional breathing synchronization induced in-phase synchronization of heart rhythms between individuals. Critically, interpersonal synchronization coincided with cardiorespiratory decoupling: respiration and heart rhythms became more out-of-phase during interaction compared to resting baselines and the unidirectional condition. Moreover, stronger interpersonal respiratory synchronization predicted greater intra-personal cardiorespiratory decoupling, particularly when participants adapted their own breathing to another person’s. These findings provide evidence that interpersonal physiological synchronization entails a trade-off with intra-personal physiological coupling, perturbing the phase relationship within one’s own physiological system. We propose that aligning one’s physiological rhythms with others strengthens self–other coupling but weakens intrapersonal coupling, pointing to a physiological mechanism of self-decoupling during social interaction.

**Graphical abstract:** When two people synchronize their breathing together, their heart rhythms also synchronize, while their intrapersonal cardiorespiratory rhythms become decoupled, with a perturbed phase-relationship between respiration and cardiac rhythms. Across two experiments, stronger interpersonal respiratory synchronization was associated with more pronounced intrapersonal cardiorespiratory decoupling. These findings suggest a physiological trade-off: aligning bodily rhythms with others strengthens self–other coupling, but may coincide with a phase shift in one’s intrinsic cardiorespiratory rhythm.

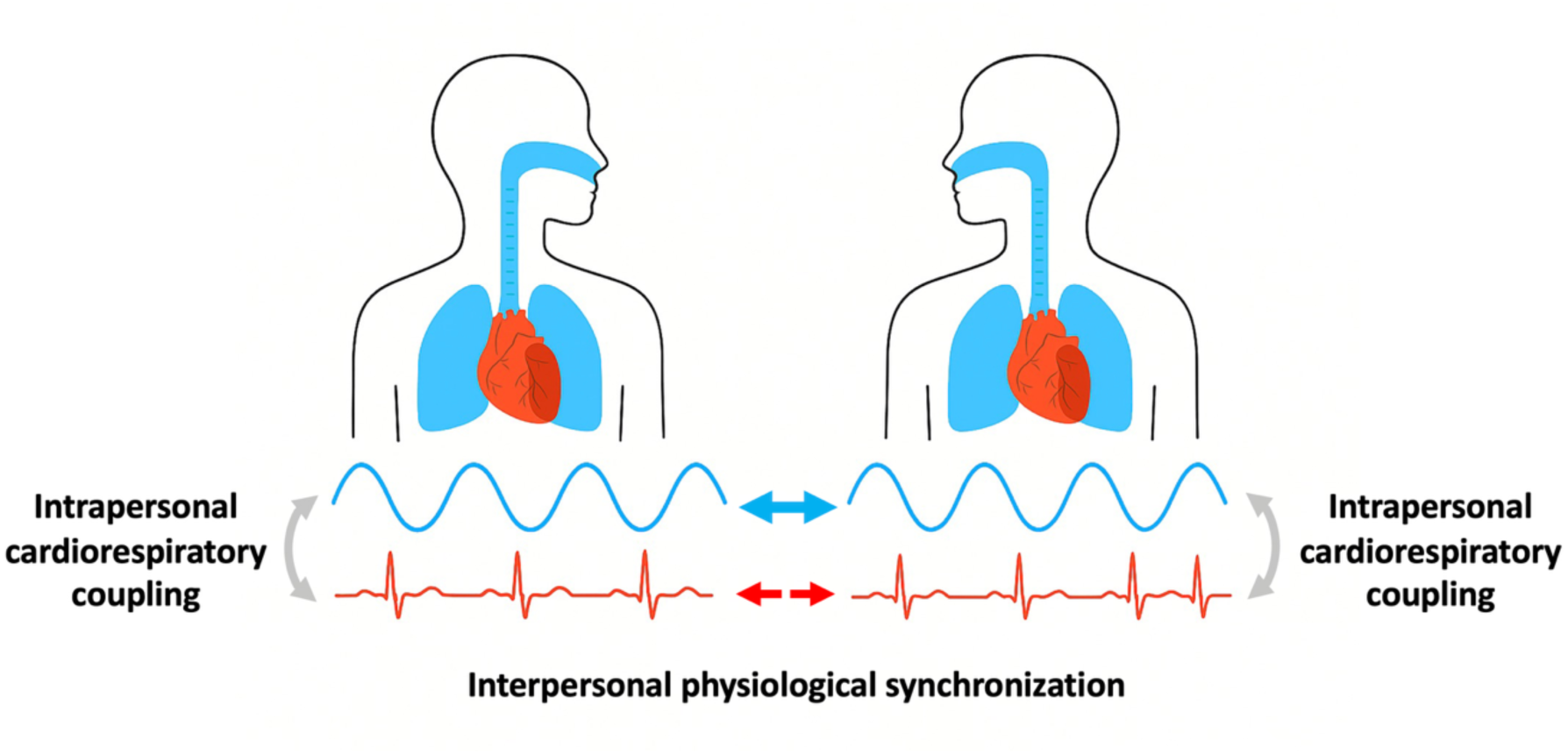

## 1 Introduction

When people engage in social interaction or simply receive visual or auditory feedback of one another, they tend to synchronize their bodily rhythms with each other both spontaneously and intentionally^1–3^. For example, people’s applause becomes synchronized following a theatre or opera performance^4^; they synchronize their walking strides when walking side-by-side^5^; and even their postures become coordinated when engaged in conversation with one another^6^. Such interpersonal synchronization has been proposed to promote self-other integration and merging^7,8^. Beyond overt motor behaviour, more recent research shows that interacting individuals also synchronize their periodic physiological signals, such as heart rhythms and respiration^9–12^.

Interpersonal physiological synchronization has been observed across a wide range of contexts^11^: mother-infant dyads show synchronized fluctuations in heart rate and respiration during face-to-face interaction^13,14^; maternal and fetal rhythms show coupling during periods of paced maternal breathing^15^; socially close dyads exhibit higher heart rate synchronization during shared fearful experiences than dyads that are not close^16^; romantic couples^17^ and attracted blind dates^18^ exhibit heart rhythm synchronization; and performers and spectators synchronize heart rhythms during collective rituals such as fire-walking, and this interpersonal synchronization is modulated by the relationship between them^9^. Behavioural coordination also modulates interpersonal physiological synchronization. For example, choir singers who sing in unison synchronize their respiration rhythms, and as a result also heart rhythms^19^; and behavioural movement synchronization has also been linked to heart rhythm synchronization^20^. Breathing rhythms also synchronize during conversation, as a consequence of the coordination between interlocutors’ utterances, even though the speakers take turns to speak and listen^21,22^. A recent study also found a causal relationship between respiratory synchronization and behavioural synchronization, which was further modulated by interpersonal differences in resting rhythms, such that partners with more similar behavioural and physiological rhythms achieved higher synchronization during interaction^23^.

Interpersonal behavioural and physiological synchronization have been proposed to follow the same dynamical principles as synchronization of other biological (e.g., fire-flies) or physical (e.g., pendulums) systems, arising from adjustments of rhythms between interacting self-sustained periodic behaviours through phase and frequency locking^24^. As a result, the underlying dynamics of human interpersonal synchronization have been widely studied using nonlinear models of coupled oscillators^17,25–27^.

However, the effects of human interpersonal synchronization extend beyond the mechanics of their coupled dynamics. Much research suggests that interpersonal behavioural and physiological synchronization lead to positive social outcomes, promoting social connection, cohesion, and prosociality^28–32^. Yet a smaller body of work indicates that such effects may not always be beneficial to the self^12^. Empirical studies have linked behavioural interpersonal synchronization to poorer self-regulation of affect^33^ and self-decoupling^27,34^, but no research has directly examined how interpersonal physiological synchronization affects one’s own intrapersonal physiological rhythms.

This question may be particularly interesting in the context of respiration, which is both under voluntary control and strongly linked to cardiac dynamics, given that respiration and heart rhythms are weakly coupled to one another^35,36^. This suggests that synchronization of breathing may influence not only self-other integration but also the coupling of one’s own internal physiological rhythms. In other words, adjusting one’s physiological rhythms to another person’s through *inter*personal physiological synchronization may affect the phase relationship in one’s own *intra*personal cardiorespiratory system, modulating the coupling between one’s respiration and heart rhythms. Understanding the consequences of interpersonal synchronization on intrapersonal “self” coupling could provide insights into dynamic blurring of self and other that underlies interpersonal coordination of behavioural and physiological rhythms.

The underlying mechanisms that modulate interpersonal synchronization are still debated, but have been linked to self and other processes. In the context of movement synchronization, as people move in and out of synchronization, they balance self-other integration and segregation, shifting attentional focus between their own and others’ actions^27,37,38^. Maintaining a clear distinction between self and other is considered crucial for joint action^39^, as it allows individuals to flexibly switch between self-other integration and segregation. Indeed, the degree to which people adapt to one another varies, with distinct interpersonal strategies emerging during coordination. For example, partners may mutually adapt^40^, one may lead while the other follows^41^, or both may maintain their own rhythm^42^, effectively decoupling from the other by focusing on their own actions. In a study of musicians performing a finger-tapping task, most participants achieved interpersonal synchronization via mutual adaptation, whereas drummers tended to ignore their partner’s timing and maintain their own tempo – yet still synchronized like two independent metronomes^42^. A mathematical model capturing these strategies revealed that mutual adaption and “follower” behaviour were characterized by strong interpersonal coupling of self-other action-perception loops and weakened within-personal coupling, reflecting dynamic self-other integration. Conversely, the non-adaptive drummer or “leader” behaviour was characterized by weak interpersonal coupling but strong intra-personal action-perception coupling, reflecting self-other segregation. This model thus suggests that strong coupling to others, leading to interpersonal synchronization, coincides with decoupling within the self.

Consistent with this prediction, Zimmermann and colleagues^34^ found neural evidence of self-decoupling during interactive finger movement coordination. When participants mutually synchronized their movements, EEG functional connectivity was reduced compared to when they improvised alone, suggesting decreased intra-brain synchronization when integrating one’s actions with those of another, potentially due to reduced self-monitoring. However, we add caution to the interpretation of this finding as the neural mechanism underlying task-based reduced intra-brain synchronization remains unclear. Conversely, being observed by the other led to a widespread increase in functional connectivity, particularly over frontal and centro-parietal regions, suggesting neural effects of heightened self-focus^34^.

Yet the functional effects of interpersonal synchronization on the self remain understudied. Theoretical work proposes that social interaction should also allow for moments of independence and disengagement from the other^43^, and that excessive social attunement, as during prolonged periods of interpersonal synchronization, may impair self-regulation^33^. Supporting this view, one study found that interpersonal movement synchrony during dyadic improvisation was associated with reduced self-regulation of affect, whereas intra-personal synchrony – measured as the coordination between one’s own limbs – opposed these effects^33^.

Taken together, these studies suggest that excessively monitoring and adapting to others’ behaviour and bodily rhythms in social settings may result in decoupled intra-personal rhythms, and consequently, poorer self-regulation. Extending these insights from motor behaviour to physiological rhythms raises new questions: does synchronizing one’s breathing with another person’s pull one away from their natural respiratory rhythms, affecting their own intra-personal cardiorespiratory synchronization? Strong interpersonal respiratory synchronization may require adjustments that weaken the stable phase relationship between heartbeats and breathing cycles observed in resting states. Investigating this trade-off between interpersonal and intra-personal coupling offers a novel way to link theories of self-other integration and segregation to physiological mechanisms, and to understand how the social self is dynamically balanced with the physiological self.

In this study, we investigate how synchronized breathing, achieved through real-time visual biofeedback of other people’s breathing rhythms, influences one’s own cardiorespiratory coordination. Previous analysis of the respiration data from this study showed that synchronization of breathing rhythms is facilitated not only by mutual interaction, but through emergence of more predictable breathing rhythms during mutual interaction^10^. Here, we analyze the effect of interpersonal breathing synchronization on interpersonal heart rhythm synchronization and intra-personal cardiorespiratory coupling. We contrast conditions of bidirectional versus unidirectional coupling, and compare cardiorespiratory coupling during interaction versus pre-interaction baselines, across two experiments. Given that respiration and heart rhythms are weakly coupled to each other, we expected to find interpersonal heart rhythm synchronization, particularly during bidirectional interaction. Building on previous theoretical work showing that mutual adaptation during interpersonal behavioural synchronization results in the increase in self-other integration but weakening of one’s own action perception loops, we expected to see decreased cardiorespiratory coupling during bidirectional interaction compared to both unidirectional interaction and baseline (i.e., breathing at rest) conditions. Furthermore, we expected cardiorespiratory coupling to decrease with increasing interpersonal synchronization of breathing rhythms, with the assumption that more self-other integration would yield more self-decoupling.

## 2 Materials & methods

### 2.1 Participants

Twenty-six participants (17 females) aged 21-35 (*M* = 24.54, *SD* = 3.39 years) took part in Experiment 1 in pairs. One pair was excluded because the participants knew each other. The remaining pairs were strangers to each other. Twenty-four participants aged 21-35 (*M* = 25.71, *SD* = 4.89 years), gender-matched to those in Experiment 1, participated in Experiment 2. All participants were healthy volunteers, with normal or corrected-to-normal vision and no history of respiratory or cardiovascular conditions. The sample size for synchronization measures was determined according to studies of interpersonal coordination at the time the study was designed^20,41^. We also ran a post-hoc power analysis based on the effect sizes from the analyses of interpersonal synchronization of breathing rhythms, reported in Konvalinka et al.^10^, which estimated sample sizes of 5-26 participants needed in order to achieve 80% power. However, given that the effect of interpersonal synchronization on cardiorespiratory coupling has not been studied before, our sample size was more exploratory for this study.

All participants gave written, informed consent prior to participation, and ethical approval was obtained from the United Ethical Review Committee for Research in Psychology, in Hungary (EPKEB).

### 2.2 Task and procedure

In experiment 1, the participants were seated in separate rooms, and did not meet each other prior to the experiment. They were each fitted with a BIOPAC wireless MP150 Bionomadix respiration belt and 3-lead ECG electrodes, and all four signals were synchronized through the BIOPAC Acknowledge software. The participants were told they would breathe together with two different people, and their instructions were to synchronize their breathing to the breathing of the other person(s). They interacted in two conditions: bidirectional condition, in which both participants received real-time visual biofeedback of the other person’s breathing signal, but not their own; and unidirectional condition, in which the participants both received real-time biofeedback of the confederate’s breathing signal, while a confederate was asked to breathe at rest and did not receive any feedback. The visual signals were presented via the Acknowledge interface. Each breathing interaction lasted 5 minutes, and a 5-minute resting baseline was recorded prior to each interaction. We thus used the baseline conditions to compare cardiorespiratory coupling – the phase relationship between one’s breathing and heart rhythm signals – during interaction (bidirectional vs. unidirectional) versus baseline (i.e., while breathing at rest). The order of the interaction conditions was counter-balanced across pairs.

Moreover, we used the data collected as part of experiment 2 in order to see if we could replicate cardiorespiratory coupling effects from experiment 1 - contrasting the interaction condition to the baseline. In experiment 2, the breathing data that emerged in the bidirectional interaction condition in experiment 1 was used as visual feedback, which was originally designed to test the effect of reciprocity versus emergence of predictable breathing patterns on interpersonal synchronization, reported in Konvalinka et al.^10^. Participants were thus recruited individually, and asked to breathe in synchrony to the signals from experiment 1 (i.e., each person interacted with one person’s breathing signal), but were told there was another participant in a different room who was producing these signals. The pre-recorded signals were resampled to the frame rate of the monitor and presented using MATLAB’s Psychtoolbox. This offline control condition lasted 5 minutes, and a 5-minute resting baseline was recorded prior to the interaction.

### 2.3 Phase analysis of heart rhythm and cardiorespiratory coupling

The respiration and ECG data were all recorded at a sampling rate of 1000 Hz. The R-peaks were extracted from each person’s signal using a peak detection algorithm, and further investigated for artefacts such as missing beats or falsely detected beats using manual detection. Any falsely detected beats were removed, and missing beats were interpolated using linear interpolation across preceding beats. In order to align the heart rate data to respiration data, the RR-intervals were converted to heart rate, resampled at 1000 Hz and interpolated using piecewise cubic spline interpolation (using interp1 function in MATLAB), and finally detrended. The respiration data were also detrended, and all the signals were normalized by computing z-scores.

The phase of each signal was estimated using the Hilbert transform, as in Konvalinka et al. ^10^. To quantify *inter*personal heart rhythm synchronization, relative phase was computed between pairs of signals. In the bidirectional interaction and baseline conditions, this was done by subtracting the phase time series from one participant from that of the other interacting participant. For the unidirectional condition, each participant’s phase signal was referenced to the confederate’s phase signal. In experiment 2, the relative phase was obtained by subtracting each participant’s phase signal from the corresponding participant’s signal in the bidirectional condition of experiment 1 (i.e., the participant whose signal they synchronized their breathing to).

For measures of *intra*personal cardiorespiratory synchronization, the resampled and interpolated heart rate signals were subtracted from the time-aligned respiration signals for each participant.

Across both heart rhythm (interpersonal) and cardiorespiratory (intrapersonal) measures, relative phase values were converted to a scale spanning -180° to +180°. The distribution of these values was then quantified by computing the percentage of occurrences within 4^°^ intervals. In the absence of synchronization, the distribution is expected to be approximately uniform across all bins. By contrast, in-phase synchronization is reflected by a concentration of values around 0^°^, anti-phase coordination by peaks near ±180^°^, and consistent phase lags by peaks at intermediate angles between -180^°^ and 0^°^ (or 0^°^ and 180^°^).

Following procedures from Konvalinka et al.^10^, we statistically compared the bidirectional (12 pairs) condition to the baseline/unidirectional (24 pairs) conditions by dividing the bidirectional recordings into two equal segments for interpersonal measures of respiratory and heart rhythm synchronization across pairs. The first segment was assigned to participant 1 and the second to participant 2, effectively yielding synchronization data across 24 pairs. Relative phase was then calculated separately for each segment, with the division corresponding to the first and second halves of the interaction period (2.5 minutes each). For the analysis of the offline control condition in experiment 2 and intrapersonal analyses of cardiorespiratory coupling, such a division was not necessary, given that we had data for 24 pairs and 24 participants, respectively.

From the relative phase distributions, we computed in-phase synchronization to quantify heart rhythm synchronization, defined as % occurrence of relative phase in the -10° to +10° window, following procedures from our previous study^10^. We also validated this range by checking whether in-phase synchronization defined as % occurrence in the -2° to 2°, -6° to +6°, and -14° to +14° windows produced consistent results. In order to quantify intrapersonal cardiorespiratory coupling, we computed the mean relative phase of the relative phase distributions between respiration and cardiac rhythms.

Finally we computed the stability of the cardiorespiratory synchronization using phase locking values (PLV), calculated between the phases of the respiration and heart rhythm signals. For each participant k, and T number of time samples, PLV was defined as:

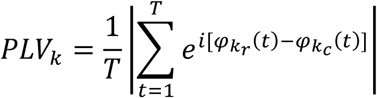

where 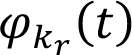 and 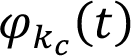 represent the instantaneous respiration and cardiac (heart rhythm) phases, respectively, at time *t*.

### 2.4 Statistical analyses

Statistical analyses were carried out using linear mixed-effects models (LMMs), with participants as random intercepts. For each LMM, we compared the full model with reduced versions, using a stepwise approach based on Akaike’s information criterion (AIC) model selection strategy. The best performing model was then evaluated using the Type III Analysis of Variance with Satterthwaite’s method. Pairwise comparisons were conducted using estimated marginal means, with p-values adjusted for multiple comparisons using Tukey’s method.

We also measured the relationship between interpersonal respiratory synchronization in the bidirectional (experiment 1) and offline control conditions (experiment 2), and intrapersonal cardiorespiratory coupling. We fit linear regression models with in-phase occurrence between people’s respiration rhythms as a predictor variable, and intrapersonal cardiorespiratory mean relative phase as a response variable.

## 3 Results

### 3.1 Bidirectional breathing synchronization led to heart rhythm synchronization

The distribution of relative phase was calculated between the two participants’ heart rhythms in the bidirectional and unidirectional interaction conditions, i.e., when participants synchronized their breathing to each other or to a confederate’s breathing, respectively, as well as in the first resting baseline condition prior to the interactions. The participants’ heart rhythms synchronized in-phase only in the bidirectional interaction condition (example data shown in Fig. 1B), while the baseline and the unidirectional condition resembled uniform relative phase distributions, as seen in Fig. 1C. This was confirmed using the Rayleigh test for uniformity, which was significant for the bidirectional interaction condition (*z =* 6.33, *p =* 0.0017), but not for the baseline (*z =* 0.88, *p =* 0.12) or the unidirectional condition (*z =* 0.52, *p =* 0.66). A linear mixed-effects model with the condition type as a fixed effect and participants as a random intercept revealed a significant main effect of condition on heart rhythm in-phase synchronization (% occurrence defined from -10° to +10°, following our previous analyses^10^), F(2,46) = 8.15, *p =* .00093, *R_c_*^2^ = 0.273, *R_m_*^2^ = 0.167. The bidirectional breathing condition showed a significant increase in in-phase synchronization compared to the baseline (β = 3.89, SE = 1.09, *t-ratio* = 3.55, *p* = .0025, *d* = 1.026) and the unidirectional breathing condition (β = 3.76, SE = 1.09, *t-ratio* = 3.44, *p* = .0036, *d* = 0.992). There was no statistical difference between the baseline and the unidirectional condition (β = -0.13, SE = 1.09, *t-ratio =* -0.12, *p* = .99, *d* = -0.034). These results were robust to other definitions of % occurrence for computing in-phase synchronization, including -2°:2° (F(2,46) = 7.42, *p =* .0016, *R_c_*^2^ = 0.31, *R_m_*^2^ = 0.144), -6°:6° (F(2,46) = 8.30, *p =* .00084, *R_c_*^2^ = 0.287, *R_m_*^2^ = 0.167), and -14°:14° (F(2,46) = 9.14, *p =* .00045, *R_c_*^2^ = 0.28, *R_m_*^2^ = 0.185) windows, with consistent pairwise relationships between the baseline, bidirectional, and unidirectional breathing conditions.

**Fig. 1.**
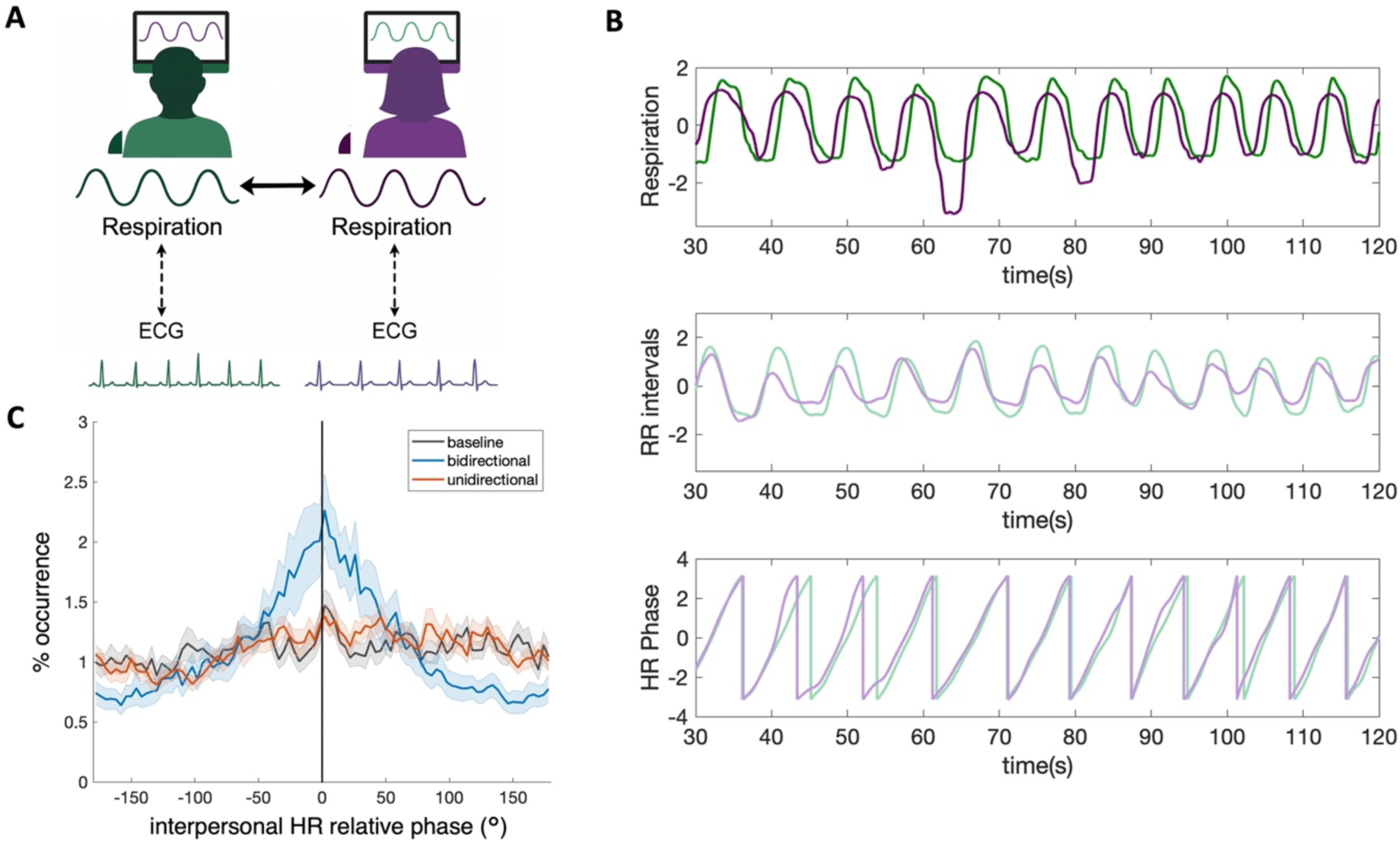
**A.** Experimental manipulation. In the bidirectional breathing condition, each participant receives real-time biofeedback of the other participant’s respiration signal in real-time. Both their respiration and ECG signals are measured, which are weakly coupled to each other. **B.** Example normalized respiration signals (top), resampled RR intervals (middle), and the phases of the resampled RR-interval signals (bottom) from two example participants in the bidirectional breathing condition. **C.** Heart rhythm synchronization: relative phase distribution between the two participants’ heart rhythm signals (i.e., resampled RR intervals) during the baseline (black), bidirectional (blue), and unidirectional (red) conditions. Shaded areas indicate standard error of the mean.

### 3.2 People’s cardiorespiratory rhythms decoupled out-of-phase during interpersonal synchronization of respiration rhythms

The distribution of relative phase was calculated between each participant’s respiration and heart rhythms during the bidirectional and unidirectional interaction conditions and the resting baseline conditions that preceded them, as shown in Fig. 2A, capturing cardiorespiratory coupling. All the conditions exhibited cardiorespiratory synchronization, which were centered at a negative relative phase difference, indicating the heart rate increases and decreases (or RR interval decreases and increases) slightly preceded inhalation and exhalation, respectively. We extracted the mean relative phase between respiratory and cardiac rhythms, as seen in Fig. 2B, in order to capture the degree of decoupling. A linear mixed-effects model with the condition (interaction vs. baseline), directionality (bidirectional vs. unidirectional), and condition x directionality interaction as fixed effects (the best performing LMM, *R_c_*^2^ = 0.436, *R_m_*^2^ = 0.211), and participants as a random intercept, revealed a significant main effect of condition on the cardiorespiratory mean relative phase F(1,69) = 25.11, *p <* .0001, and a significant condition x directionality interaction F(1,69) = 8.11, *p =* .0058, but no effect of directionality F(1,69) = 2.38, *p =* .13. Pairwise comparisons revealed a significant decrease in the cardiorespiratory mean relative phase during the bidirectional interaction condition compared to the preceding baseline (β = -20.05, SE = 3.61, *t-ratio = -*5.56, *p* < .0001, *d* = -1.604), as well as compared to the unidirectional interaction (β = -11.20, SE = 3.61, *t-ratio =* -3.11, *p* = .014, *d* = -0.896). There was no difference in the cardiorespiratory mean relative phase between the unidirectional interaction and the preceding baseline (β = -5.52, SE = 3.61, *t-ratio =* 1.53, *p* = .43, *d* = -0.442).

**Fig. 2.**
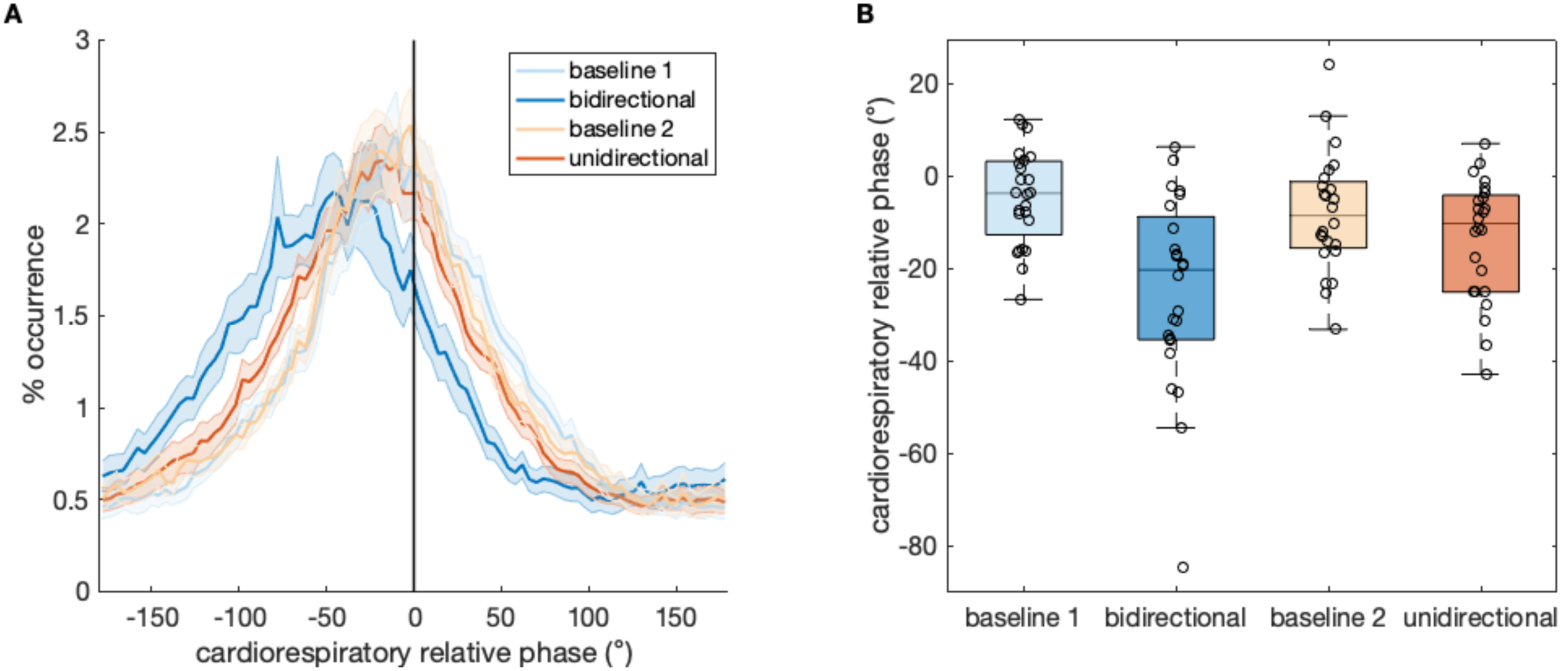
Cardiorespiratory coupling in Experiment 1, showing effects of interpersonal synchronization of respiration rhythms on intra-personal synchronization of cardiorespiratory rhythms. **A.** Distribution of relative phase (from -180° to 180°) across the condition types (baseline vs. interaction) and directionality (bidirectional vs. unidirectional). The baselines correspond to the resting baseline recorded prior to each of the interaction conditions, where baseline 1 preceded the bidirectional interaction, while baseline 2 preceded the unidirectional interaction (baseline numbers do not correspond to the order of conditions, which is counterbalanced). Shaded areas indicate standard error of the mean. **B.** Box plots of the cardiorespiratory mean relative phase across the four conditions (baseline 1, bidirectional interaction, baseline 2, unidirectional interaction).

To test whether the decoupling could be explained by the variations in the breathing frequency, which showed high variability as reported in Konvalinka et al.^10^, we added the breathing frequency as a covariate to the model, both as a fixed effect on its own as well as in interaction with the condition and directionality. However, there was no statistical difference in model comparisons between the original simpler model (*LMM1*: meanrelphase ∼ Condition * Directionality + (1|Subject)) and the more complex models with frequency as a fixed effect (*LMM2*: meanrelphase ∼ Condition * Directionality + Frequency + (1|Subject), *p =* 0.59; *LMM3*: meanrelphase ∼ Condition * Directionality * Frequency + (1|Subject), *p =* 0.34). Hence, the effect of bidirectional interaction on cardiorespiratory coupling could not be explained by the breathing frequency.

To test whether the interaction had an effect on the stability of the cardiorespiratory phase relationship, we tested the effect of condition, directionality, and condition x directionality interaction on the PLV between the respiration and cardiac rhythms. Model comparisons revealed the simplest model, with only the condition as a fixed effect and participants as a random intercept, to be the best performing model. The condition (interaction vs. baseline) did not have a significant effect on the PLVs, F(1,71) = 0.05, *p =* .83, *R_c_*^2^ = 0.302, *R_m_*^2^ = 0.

To investigate whether the cardiorespiratory decoupling effect would be replicated with another group of participants, who synchronized their respiration to the respiration signals that emerged in the bidirectional condition in Experiment 1, we tested the effect of the condition (interaction vs. baseline) on the mean relative phase in Experiment 2. Fig. 3A suggests that people’s cardiorespiratory rhythms also decoupled during interaction in this scenario, showing a peak in the distribution of relative phase in the negative direction once again. The relative phase distribution for the baseline condition was centered closer to 0 degrees, suggesting in-phase cardiorespiratory synchronization during the baseline. A linear mixed-effects model with the condition as the fixed effect and participants as random intercepts, showed that synchronizing one’s own breathing to another person’s (from the bidirectional condition) decreased the mean relative phase in comparison to the resting baseline (Fig. 3B), F(1,23) = 5.75, *p =* .025, *R_c_*^2^ = 0.211, *R_m_*^2^ = 0.097, showing evidence of cardiorespiratory decoupling. This effect could not be explained by the breathing frequency alone, as shown through model comparisons when adding frequency as a covariate or fixed effect in the linear mixed-effects model.

**Fig. 3.**
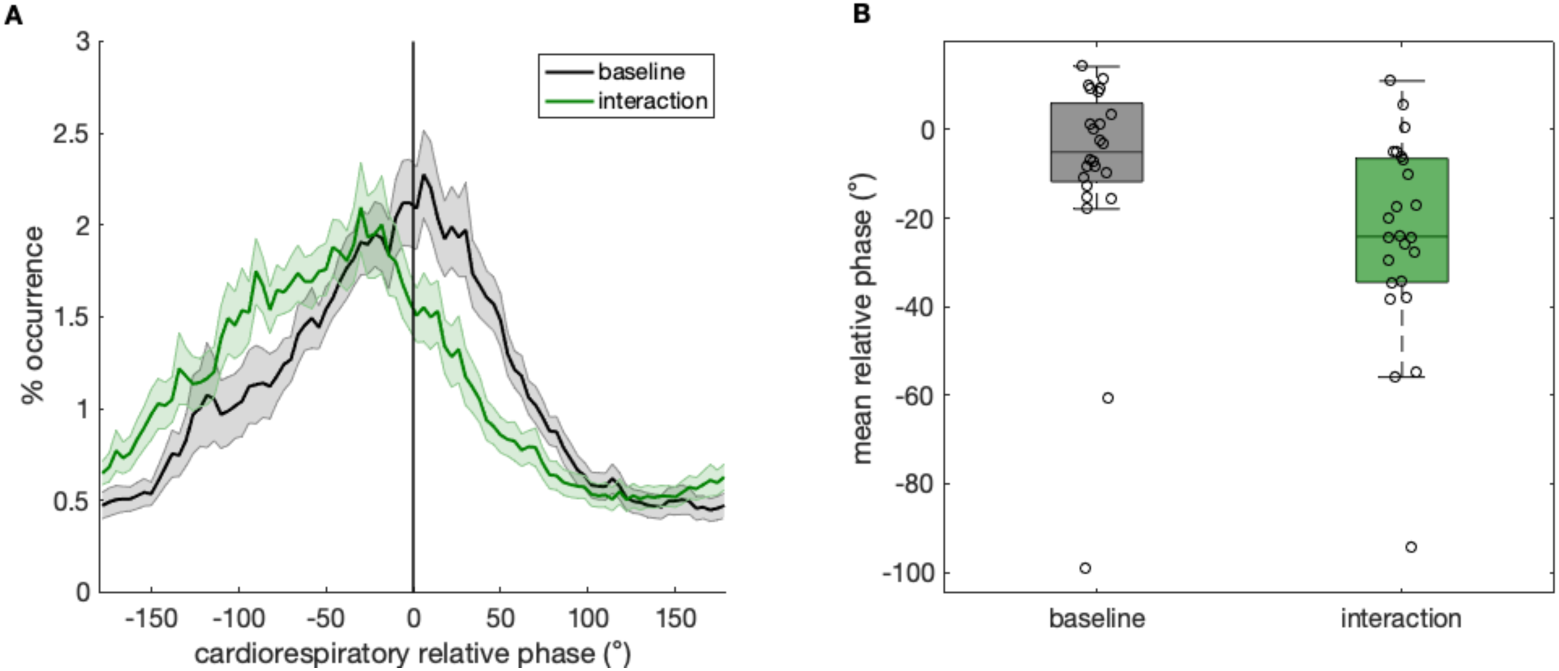
Cardiorespiratory coupling in Experiment 2. **A.** Distribution of relative phase (from - 180° to 180°) across the resting baseline and interaction (offline control) conditions. Shaded areas indicate standard error of the mean. **B.** Box plots of the mean relative phase during baseline and interaction conditions across all participants.

### 3.3 Higher *inter*personal synchronization of respiration rhythms was associated with greater *intra*personal cardiorespiratory decoupling

In order to test the relationship between interpersonal synchronization of respiration rhythms and intrapersonal cardiorespiratory (de)coupling, we fitted linear regression models relating the in-phase respiratory synchronization of the paired participants’ respiration rhythms to individual cardiorespiratory mean relative phase, in both the bidirectional interaction condition (experiment 1) and the offline control condition (experiment 2).

In the bidirectional interaction condition (Fig. 4A), the more in-phase synchronized the participants were (% occurrence defined from -10° to +10°), the more out-of-phase their cardiorespiratory rhythms were (*r*^2^ *=* 0.224, *p =* 0.0194). This relationship was robust to different definitions of in-phase synchronization % occurrence, including -2°:2° (*r*^2^ *=* 0.213, *p =* 0.0231), -6°:6° (*r*^2^ *=* 0.225, *p =* 0.0193), and -14°:14° (*r*^2^ *=* 0.223, *p =* 0.0197). However, it was not robust to removal of an individual with an extreme cardiorespiratory phase value (−94.5°), after which the relationship was no longer significant (−10°:10° % occurrence: *r*^2^ *=* 0.11, *p =* 0.12).

**Fig. 4.**
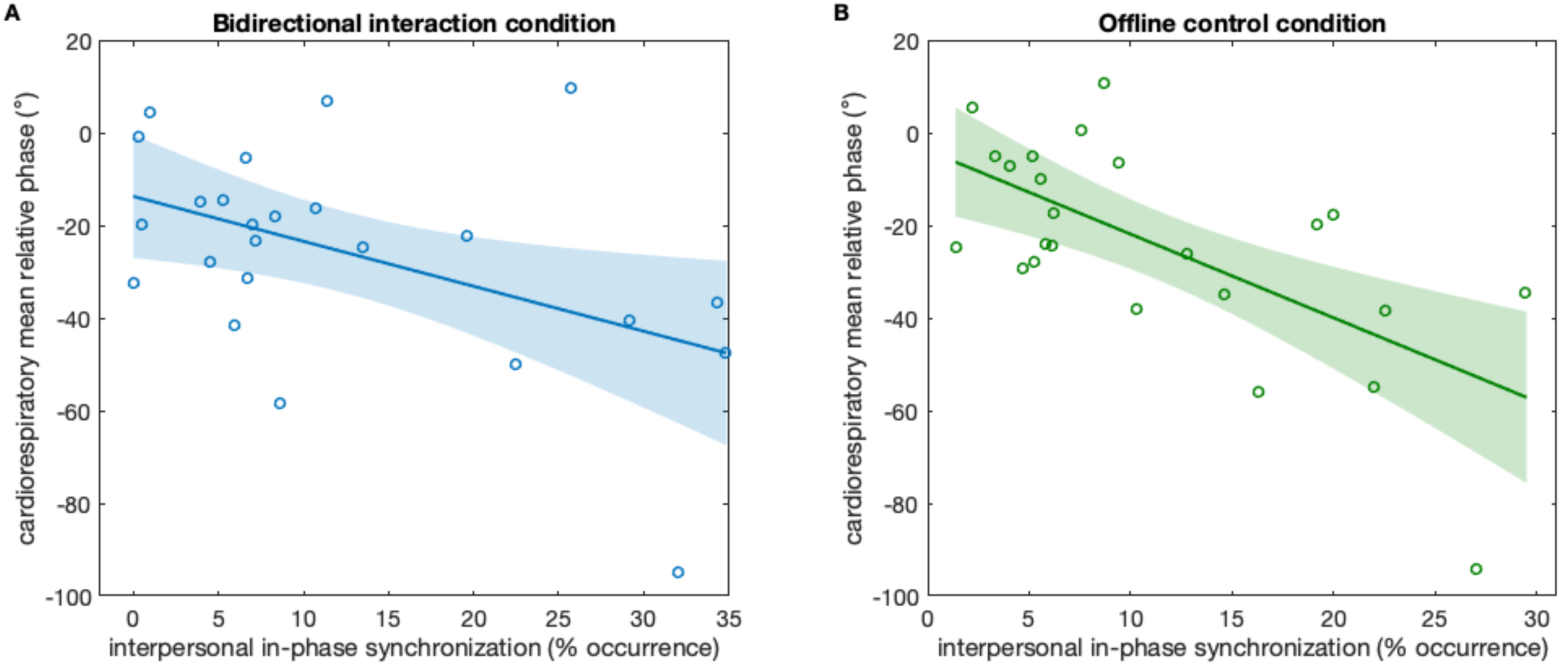
Relationship between interpersonal synchronization of respiration rhythms (in-phase % occurrence, from -10° to +10°) and intrapersonal cardiorespiratory decoupling (cardiorespiratory mean relative phase) during the **A.** bidirectional interaction condition (experiment 1) and **B.** offline control condition (experiment 2). Shaded areas indicate 95% confidence intervals.

In the offline control condition (Fig. 4B), the same relationship between interpersonal synchronization and intrapersonal cardiorespiratory decoupling was observed (*r*^2^ *=* 0.43, *p =* 0.0005). Importantly, this association remained significant after removal of one individual with an extreme cardiorespiratory phase value (–94.3 degrees), *r*^2^ *=* 0.31, *p =* 0.0057. In this condition, intrapersonal cardiorespiratory coupling was also more directly related to the extent to which individuals adapted their breathing to the visually presented breathing signal.

## 4 Discussion

We investigated how synchronizing breathing with another person modulates interpersonal heart rhythm synchronization and intra-personal cardiorespiratory coupling. Given that breathing is weakly coupled to heart rate^35,36^, such that heart rate increases during breathing inhalation and decreases during exhalation, we expected to find interpersonal heart rhythm synchronization when two people synchronized their breathing together. As expected, during bidirectional interaction, when people reciprocally synchronized their breathing together, interpersonal heart rhythm synchronization emerged. Heart rhythm synchronization did not emerge in the unidirectional interaction, between participants and a confederate. Interestingly, interpersonal synchronization of physiological rhythms led to a decoupling of intrapersonal cardiorespiratory rhythms, with a larger out-of-phase relationship between respiration and heart rate during bidirectional interaction compared to breathing at rest (or compared to synchronizing one’s breathing to a confederate’s breathing in the unidirectional interaction). Moreover, the more people interpersonally synchronized their breathing with another person, the more decoupled their intrapersonal cardiorespiratory rhythms became. This was a robust effect in experiment 2, where participants synchronized their breathing rhythms unidirectionally to the breathing rhythms that emerged from reciprocal interaction in experiment 1. The effect was also significant in experiment 1, where participants bidirectionally synchronized their breathing together, but it was not robust to removal of one extreme value of cardiorespiratory decoupling. Taken together, our results indicate that interpersonal physiological synchronization leads to a decoupling of one’s own physiological rhythms.

This study provides empirical evidence that synchronizing one’s own bodily rhythms with another person, which requires self-other integration, results in a self-decoupling within one’s own physiological system. It also suggests that synchronized breathing pulls people away from their own natural physiological rhythms, which may weaken the stable phase relationship within their own cardiorespiratory system, i.e., between their breathing and heart rhythm cycles. The more one synchronizes with another, the more one is presumably pulled away from their own natural rhythms, which may explain the linear association between the amount of interpersonal synchronization and intrapersonal decoupling, as robustly shown in experiment 2. This empirical evidence builds on previous neural evidence of self-decoupling^34^, as well as theoretical findings within computational modeling of interpersonal behavioural synchronization. Such models have shown that interpersonal action coordination where one person adapts their behaviour to another’s leads to a strengthening in between-personal action-perception loops but a weakening in one’s own within-personal action-perception coupling^27,44^.

While the linear association between bidirectional interpersonal synchronization and intrapersonal cardiorespiratory decoupling was not robust to extreme values in experiment 1, we caution that the comparison in experiment 1 involves relating a dyadic (interpersonal) measure to an individual (intrapersonal) measure. Because respiratory synchronization in a bidirectional interaction emerges from mutual adaptation, it is unclear whether both partners contributed equally to the observed synchronization or whether it was primarily driven by one individual. As a result, attributing the dyadic synchronization measure to individual-level cardiorespiratory dynamics is not straightforward. In contrast, this concern does not apply to the offline control condition, where respiratory adaptation was unidirectional (participant adapting to an externally presented breathing signal), allowing interpersonal synchronization to be more directly interpreted as reflecting individual regulatory adjustments.

It could be argued that the frequency of breathing plays a role in the cardiorespiratory phase relationship, as shown in previous work^45–47^, such that e.g., slower breathing may result in a larger out-of-phase relationship between respiration and heart rate. However, the breathing frequency did not affect the cardiorespiratory phase relationship, or mediate the effect of interpersonal synchronization on cardiorespiratory coupling. Ideally, we could have better disassociated the effect of interpersonal synchronization and the breathing frequency by having a wider range of produced breathing frequencies that people had to synchronize to. However, due to our small sample size and uncontrolled breathing frequencies, this is a limitation of this work. In addition to the breathing frequency, tidal volume, respiration depth, and vagal tone have been shown to influence the magnitude of respiratory sinus arrhythmia and cardiorespiratory coupling^48–50^. We did not obtain tidal volume measurements in this study; however, tidal volume and vagal tone are primarily known to modulate cardiorespiratory coupling strength, whereas their influence on phase offset is less well understood. We considered including vagal tone as a covariate, but this measure is confounded by the breathing frequency ^49^, which varied substantially across participants. Including it would thus make it difficult to disentangle the effects of vagal tone from those of breathing frequency. Future studies should better control for these physiological variables and their potential effects on cardiorespiratory phase dynamics.

Interestingly, even the baseline conditions in experiment 1 showed a slight out-of-phase cardiorespiratory relationship, while the baseline condition in experiment 2 did not. This may be explained by carryover effects from the interaction conditions, as the order of bidirectional and unidirectional conditions was counterbalanced across pairs. Half of the baselines preceding each condition were recorded after one of the interaction conditions. For example, if one person started with the unidirectional interaction condition, the baseline preceding the bidirectional condition was recorded after the unidirectional interaction and before the bidirectional one. Notably, in experiment 2, all the baselines were recorded first, as participants only engaged in one interaction. However, the extent to which self-decoupling carries over after interpersonal synchronization remains to be investigated in future research.

This research also builds on previous work by Galbusera and colleagues^33^ showing that interpersonal synchronization impedes self-regulation of affect. Similarly, previous research suggests that weakened self-regulation may modulate interpersonal synchrony. For example, anxious parents have been reported to have higher physiological synchrony with their infants than less anxious parents, due to greater self-reactivity to small-scale fluctuations in their infants’ arousal^51^. In other words, when their infants experience increased arousal, instead of downregulating, the anxious parents experience increased arousal as well, which may reflect poorer self-regulation.

The question remains whether the physiological self-decoupling we observe here is indicative of weakened self-regulation, or is merely a physiological correlate that might relate to self-regulatory processes. In addition, what are the behavioural or psychological consequences of such self-decoupling within the physiological system? Are we less able to regulate our arousal to different situations when we have synchronized our physiological rhythms with other people’s rhythms? Is this mechanism related to self-monitoring, such that decreased self-monitoring, as when integrating oneself with others in coordination tasks, leads to physiological self-decoupling and less self-regulation? Unfortunately, as we did not include any subjective measures of self-regulation in this study, we cannot determine whether the altered cardiorespiratory phase relationship corresponds to changes in self-monitoring or regulation of affect or arousal.

Furthermore, other factors may have influenced participants’ degree of adaptation to the biofeedback signals, and thereby their cardiorespiratory coupling, including attention^52,53^, engagement^54^, or interoceptive awareness. Previous work has shown that increasing interoceptive focus, such as focusing attention to one’s own breath during social interaction, leads to higher interpersonal physiological^55^ and neural synchronization^56^. In contrast, our task required participants to adapt their breathing to an external signal, potentially shifting attention away from internal monitoring and toward the other. Hence, we would expect higher interoceptive focus to lead to less adaptation, and hence lower interpersonal physiological synchronization. However, as we did not manipulate interoceptive focus or measure interoceptive awareness, we cannot determine how individual differences in bodily awareness may have modulated responsiveness to the feedback. Whether interpersonal synchronization is facilitated by increased attention to others, or individual differences in interceptive awareness thus remains to be explored.

Finally, as our participants did not engage in a face-to-face interaction, but were merely coupled through biofeedback of each other’s physiological rhythms, it remains unclear how synchronization of physiological rhythms was subjectively experienced. Did this low-level mutual adjustment of physiological rhythms evoke stronger feelings of togetherness^57^ and affiliation^7^ or reflect joint attention and engagement^54,58^? In our previous analysis of this study, we did not find any association between breathing synchronization and the perceived amount of synchronization or any social bonding measures^10^, suggesting that the participants did not monitor the degree of coupling between them or experience more affiliation or self-other overlap when they synchronized more. Nevertheless, the task required reciprocal exchange of physiological signals, even if it was not through face-to-face interaction. It is therefore possible that mutual adjustment of physiological rhythms primarily affected internal regulatory dynamics rather than shared subjective experience.

Importantly, the structured and remote nature of the paradigm limits the extent to which these findings can be generalized to more naturalistic, face-to-face social interactions. In everyday interactions, physiological synchronization accompanies spontaneous behaviour, multimodal sensory cues, physical proximity^54^, and mutual adaptation, which may modulate the relationship between interpersonal and intrapersonal coupling. Future work should therefore examine whether similar cardiorespiratory phase shifts emerge during naturalistic face-to-face interactions. Additionally, combining intra- and interpersonal physiological synchronization measures with subjective measures of arousal, engagement, and self-regulation will help determine the psychological significance of cardiorespiratory decoupling.

## Conclusion

The present study is the first to show intrapersonal physiological effects of interpersonal synchronization, suggesting that when we synchronize our physiological rhythms with others, our own within-personal physiological rhythms become decoupled through an altered phase relationship. This is in line with previous research showing decoupling of one’s own action-perception loops during action synchronization, and in line with previous proposals that the mechanism underlying interpersonal synchronization may involve self and other blurring and merging. Future work could investigate how physiological decoupling during interpersonal synchronization impacts self-regulation and self-other blurring, and more directly, what consequences such decoupling has for the self.

## Acknowledgements

This work is supported by the Carlsberg Foundation’s Semper Ardens: Accelerate grant, “Self, other, and we: mechanisms and dynamics of self-other integration in social observation an interaction”, CF22-1251.

## Author contributions

IK: conceptualization, funding acquisition, project administration, formal analysis, data curation, software, visualization, writing – original draft, writing – review & editing. NS: conceptualization, visualization, resources, supervision, writing – review & editing. GK: conceptualization, visualization, resources, supervision, writing – review & editing.

